# ARSENAL: Learning Transferable Regulatory DNA Representations with Targeted Short-Context Language Models

**DOI:** 10.64898/2026.02.05.703637

**Authors:** Aman Patel, Anshul Kundaje

## Abstract

DNA language models (DNALMs) aim to learn representations of genomic sequence for variant interpretation, regulatory prediction, and sequence design. Most DNALMs are trained on whole genomes and long contexts, but regulatory DNA poses a distinct challenge: functional elements are sparse, context dependent, and encoded by short transcription factor motif syntax embedded in extensive background sequence. We introduce **ARSENAL**, a short-context masked DNA language model pretrained on ENCODE candidate cis-regulatory elements. ARSENAL recovers diverse transcription factor motifs *de novo* and improves zero-shot regulatory variant effect prediction relative to other DNALM foundation models. ARSENAL embeddings also improve supervised regulatory sequence models at predicting chromatin accessibility and regulatory variant scoring. Finally, ARSENAL serves as an efficient generative prior, enabling multi-objective regulatory sequence design with supervised oracles. ARSENAL shows that targeted self-supervised pretraining on regulatory regions can learn biologically meaningful and transferable regulatory representations without genome-scale training, long contexts or task-specific labels.

Code is available at https://github.com/kundajelab/regulatory_lm. Models and data are shared at https://sageb.io/ydjhqM

## 1 Introduction

DNA language models (DNALMs) aim to learn general-purpose representations of genomic sequence for variant interpretation, regulatory prediction, and sequence design. Motivated by the success of protein language models [31, 19, 17], recent DNALMs have scaled masked and autoregressive architectures to long sequence contexts and trained them on one or more whole genomes. Representative examples include Nucleotide Transformer, HyenaDNA, DNABERT-2, Caduceus, GENA-LM, Evo, and Evo2 [26, 36, 13, 23, 2, 11, 7, 28, 8]. These models seek to learn broadly useful sequence representations by increasing model capacity, training corpus size, and context length.

Regulatory genomics presents a distinct challenge for this paradigm. The human genome contains diverse functional sequence classes embedded within a large fraction of neutrally evolving background DNA. Proteincoding regions comprise only about 1.5% of the genome, while non-coding *cis*-regulatory elements are estimated to span roughly 5–20% and control when and where genes are expressed [9]. These regulatory elements, including promoters and enhancers, encode short transcription factor binding sites whose affinity, spacing, orientation, and combinatorial organization shape regulatory output.

This syntax is difficult to learn from genome-wide self-supervision. In contrast to the dense and regular constraints of coding sequence, regulatory syntax is sparse, heterogeneous, combinatorial, and context dependent [25]. Functional motif-scale patterns are rare relative to background composition and repetitive sequence, so genome-wide language modeling objectives may emphasize non-functional statistics over regulatory grammar. Consistent with this concern, recent benchmarks have shown that whole-genome DNALMs often perform poorly on regulatory tasks such as variant effect prediction and regulatory activity inference, and may fail to capture nucleotide co-dependencies characteristic of functional motifs [25, 30, 20, 6, 32]. These findings suggest that scaling context length and corpus size alone does not guarantee learning of motif-scale regulatory syntax.

Several approaches attempt to inject biological structure into DNALMs. For example, GPN-style models use evolutionary conservation and multiple sequence alignments to improve variant effect prediction [6, 35]. These approaches are powerful, but alignment-based formulations may be less flexible when alignments are unavailable and can complicate generation or synthetic design. This motivates complementary strategies that bias learning toward regulatory function while preserving standard single-sequence inference and generation.

Here we introduce ARSENAL (**A R**egulatory **SE**quence **N**ucleic **A**cid **L**earner), a short-context masked DNALM pretrained on ENCODE candidate *cis*-regulatory elements (cCREs), a high-confidence set of human regulatory elements identified across diverse cellular contexts [10, 21]. ARSENAL tests a simple hypothesis: targeting self-supervised pretraining to functionally enriched regulatory regions can learn biologically meaningful and transferable representations of regulatory DNA without genome-scale contexts or task-specific labels. We show that ARSENAL recovers diverse transcription factor motifs *de novo*, captures interpretable motif-scale nucleotide dependencies, improves zero-shot regulatory variant effect prediction relative to prior single-sequence DNALMs, provides embeddings that improve supervised chromatin accessibility prediction and regulatory variant scoring, and supports targeted regulatory sequence design. Together, these results establish targeted short-context pretraining as a practical strategy for learning useful representations of regulatory DNA.

## 2 Results

### 2.1 The ARSENAL model: targeted pretraining for regulatory DNA

We introduce **ARSENAL**, a masked DNA language model designed to learn local regulatory sequence syntax from functionally enriched genomic regions (Fig. 1). ARSENAL is pretrained on approximately 2.3 million ENCODE candidate *cis*-regulatory elements (cCREs), which include promoters, enhancers, CTCF-bound elements, and other regulatory loci [10, 21]. Compared to whole-genome pretraining, this targeted corpus increases the density of regulatory motif signal relative to background sequence.

**Figure 1:**
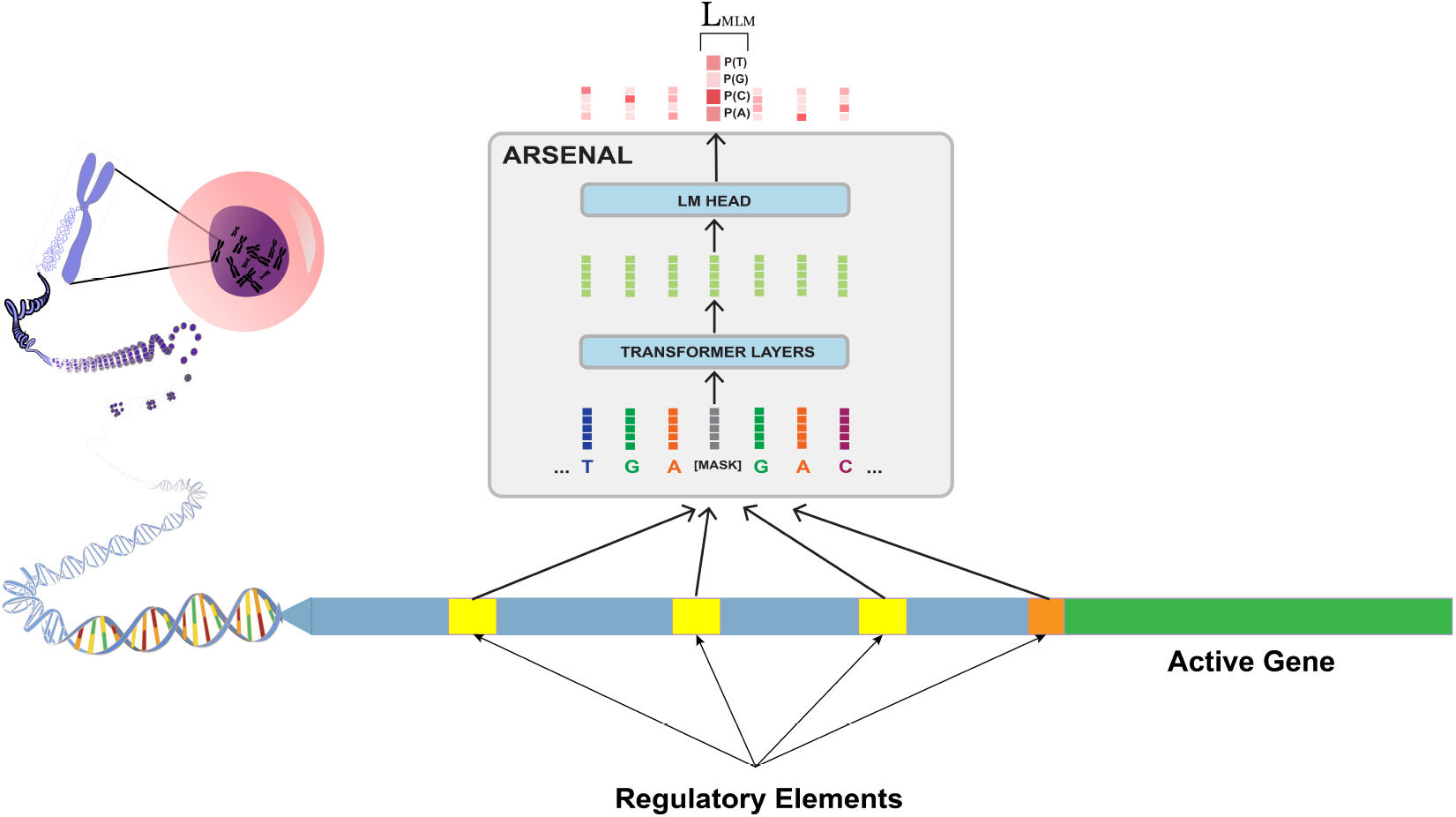
Overview of ARSENAL. ARSENAL is a short-context masked DNA language model pretrained on candidate *cis*-regulatory elements (cCREs). [1]

**Figure 2:**
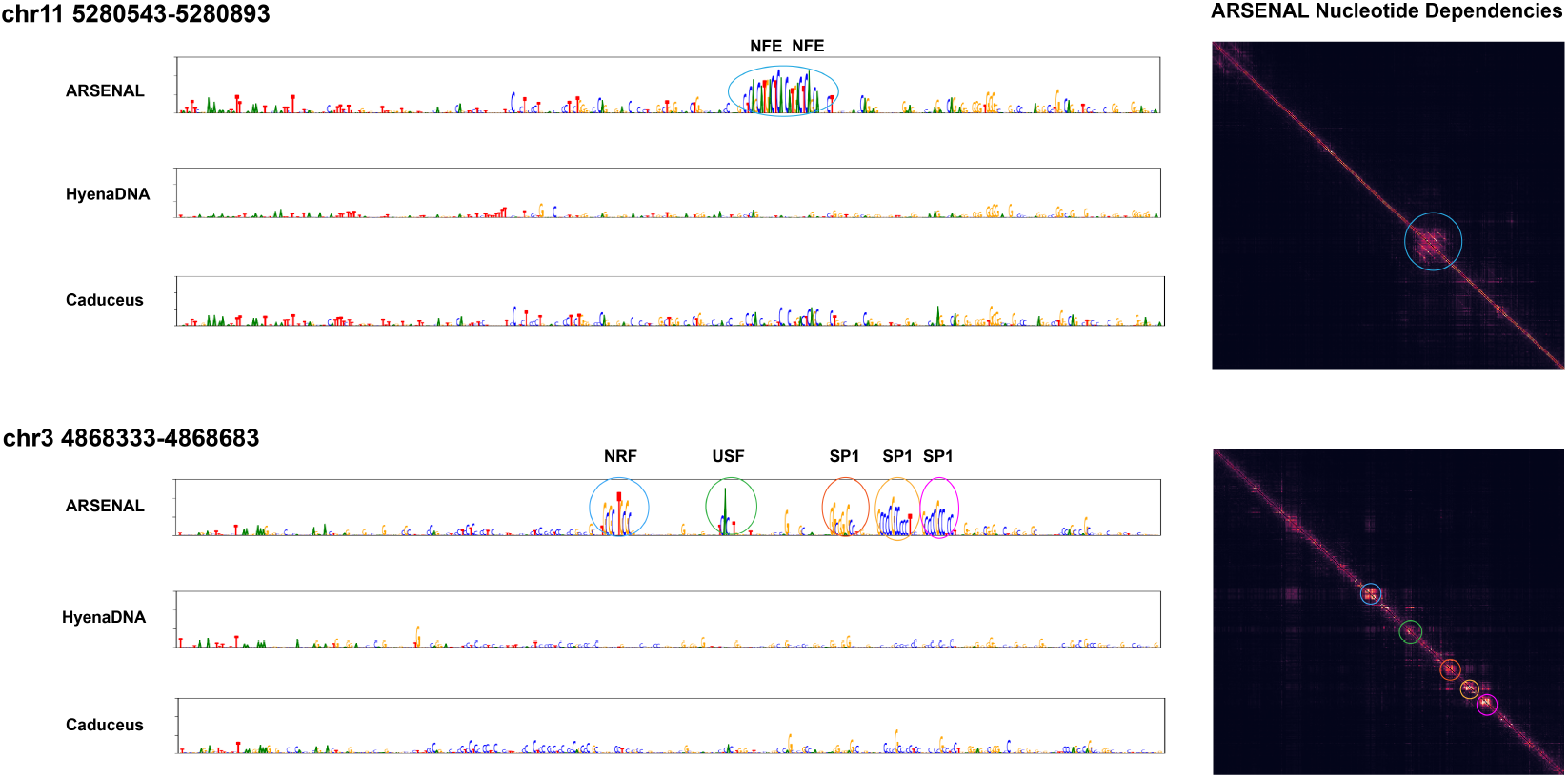
Zero-shot likelihood reconstructions and nucleotide dependency maps at two representative regulatory loci. Left: per-base likelihood reconstructions from ARSENAL, HyenaDNA, and Caduceus for an enhancer of the *β*-globin gene on chromosome 11 and a promoter-like cCRE on chromosome 3. Right: ARSENAL nucleotide dependency maps for the same loci. Circled regions in the dependency maps correspond to motif-scale features highlighted in the likelihood tracks.

ARSENAL operates on 350 bp sequence contexts, matching the local scale at which many transcription factor motifs and motif combinations occur. The model is trained with a standard masked language modeling objective: a subset of input tokens is masked, perturbed, or left unchanged, and the model is trained to predict the original nucleotide from surrounding context. This objective produces position-specific nucleotide likelihoods that can be directly interrogated for motif discovery and used for zero-shot variant scoring by comparing reference and alternate allele likelihoods. To reduce learning from repetitive sequence features that can dominate likelihood-based models, we exclude masked-token loss contributions from long soft-masked repeat stretches during pretraining (Methods).

### 2.2 ARSENAL recovers regulatory motif syntax from zero-shot likelihoods

An informative test of a regulatory DNALM is whether its zero-shot likelihoods recover known motif-scale functional features: nucleotides within transcription factor binding sites should be predicted with structured, high-confidence likelihoods, while surrounding background sequence should contribute less coherent signal. We evaluate this property in two complementary ways: locus-level reconstructions and nucleotide dependency maps, and large-scale motif identification and TF-MoDISco analyses across regulatory regions.

First, we visualize per-base likelihood reconstructions at representative loci and compare them to other DNALMs. These likelihoods are computed by successively masking each position and predicting the probability of the true nucleotide given the remaining sequence context. At an enhancer of the *β*-globin gene on chromosome 11, ARSENAL recovers two evolutionarily conserved MAFK/NFE motifs previously reported at this locus. At a promoter-like cCRE on chromosome 3, ARSENAL recovers multiple SP1, USF, and NRF family motifs [22]. In contrast, HyenaDNA and Caduceus—the two single-nucleotide-tokenized models evaluated in the DART-EVAL DNALM benchmark—produce substantially weaker or less structured reconstructions at these loci.

To test whether ARSENAL captures motif-level *dependencies* rather than only per-position confidence, we compute nucleotide dependency maps, which quantify how predictions at one position depend on nucleotides at other positions [32]. ARSENAL shows clear diagonal dependency blocks aligned to recovered motifs, consistent with learning motif-scale syntax. We also observe weaker off-diagonal blocks linking distinct motifs, suggesting that the model may capture motif co-occurrence patterns in addition to individual motif structure.

Second, we quantify the breadth of learned motif syntax using the DART-EVAL motif identification benchmark, which measures how often known binding motifs receive higher likelihood than shuffled versions of the same motifs [25]. ARSENAL outperforms other DNALMs on this benchmark (Fig. 3A), indicating broad recognition of regulatory motifs in a zero-shot setting.

**Figure 3:**
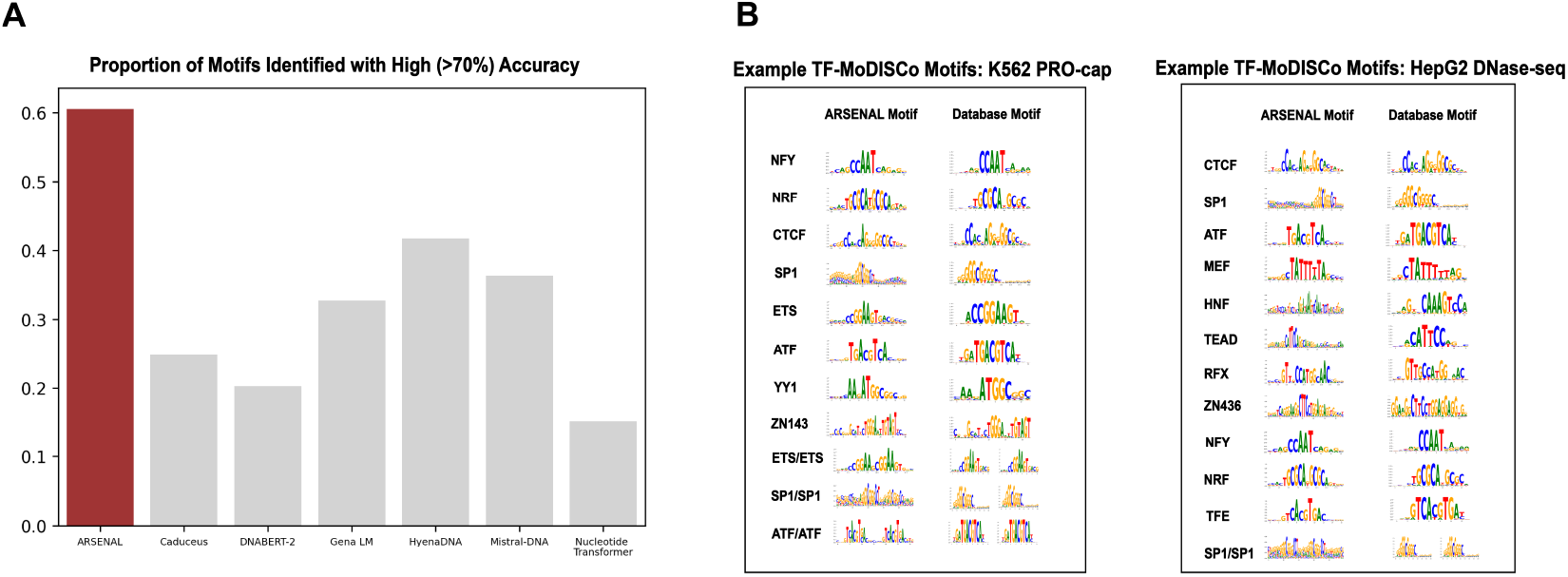
Large-scale zero-shot motif discovery with ARSENAL. (A) DART-EVAL motif identification benchmark comparing ARSENAL with other DNALMs. (B) Representative TF-MoDISco patterns discovered from ARSENAL likelihood-based contribution scores in K562 PRO-cap peaks and HepG2 DNase-seq peaks, matched to canonical motifs from HOCOMOCO [34].

As a complementary analysis, we apply TF-MoDISco to likelihood-based contribution scores across large sets of regulatory sequences [29]. In K562 promoter-proximal regions associated with nascent transcription, ARSENAL recovers canonical promoter-associated motifs including NRF, NFY, ETS, and others (Fig. 3B). In HepG2 DNase-seq peaks, which include a broader mixture of enhancers and promoters, ARSENAL recovers a cell-type specific motif repertoire, including HNF, TEAD, and MEF2 family motifs alongside more general regulatory motifs. Together, these analyses show that ARSENAL learns biologically meaningful regulatory motif syntax from sequence-only self-supervision and provides interpretable likelihood and dependency maps for mechanistic inspection of individual loci.

### 2.3 ARSENAL improves zero-shot variant effect prediction in regulatory QTL benchmarks

We next asked whether ARSENAL’s motif-level likelihood structure translates into better zero-shot variant effect prediction, a central application of regulatory sequence models. Beyond its practical importance for prioritizing non-coding variants, this task provides a stringent test of counterfactual generalization and calibration: the model must assign quantitatively meaningful probabilities at single-nucleotide resolution such that a single base change produces an appropriate shift in likelihood. This sensitivity must emerge from sequence learning alone, since ARSENAL is not trained on variant annotations or allele-specific measurements.

We evaluate ARSENAL on the DART-EVAL zero-shot variant scoring benchmark [25]. For masked language models, single-nucleotide variant effects are computed by masking the variant position and comparing the probability assigned to the alternate allele with the probability assigned to the reference allele. This likelihood-ratio score is biologically intuitive for regulatory DNA because high-effect non-coding variants often create, disrupt, or weaken transcription factor binding motifs.

We consider two DART-EVAL regulatory QTL datasets in lymphoblastoid cell lines: dsQTLs measuring effects on DNase-seq chromatin accessibility and caQTLs measuring effects on ATAC-seq accessibility [25]. These variants were derived from African-ancestry population cohorts, providing a realistic distribution of common regulatory variation. Following DART-EVAL and ChromBPNet conventions, correlations are evaluated on variants with significant measured effects (Methods). Across both dsQTL and caQTL settings, ARSENAL achieves higher correlation with experimental effect sizes than other DNALMs evaluated in the same framework (Fig. 4). Notably, ARSENAL uses only 350 bp of sequence context around each variant, whereas the original DART-EVAL evaluation uses 2,114 bp windows, suggesting that accurate local regulatory variant scoring can be achieved with short contexts when motif-scale syntax is learned effectively.

**Figure 4:**
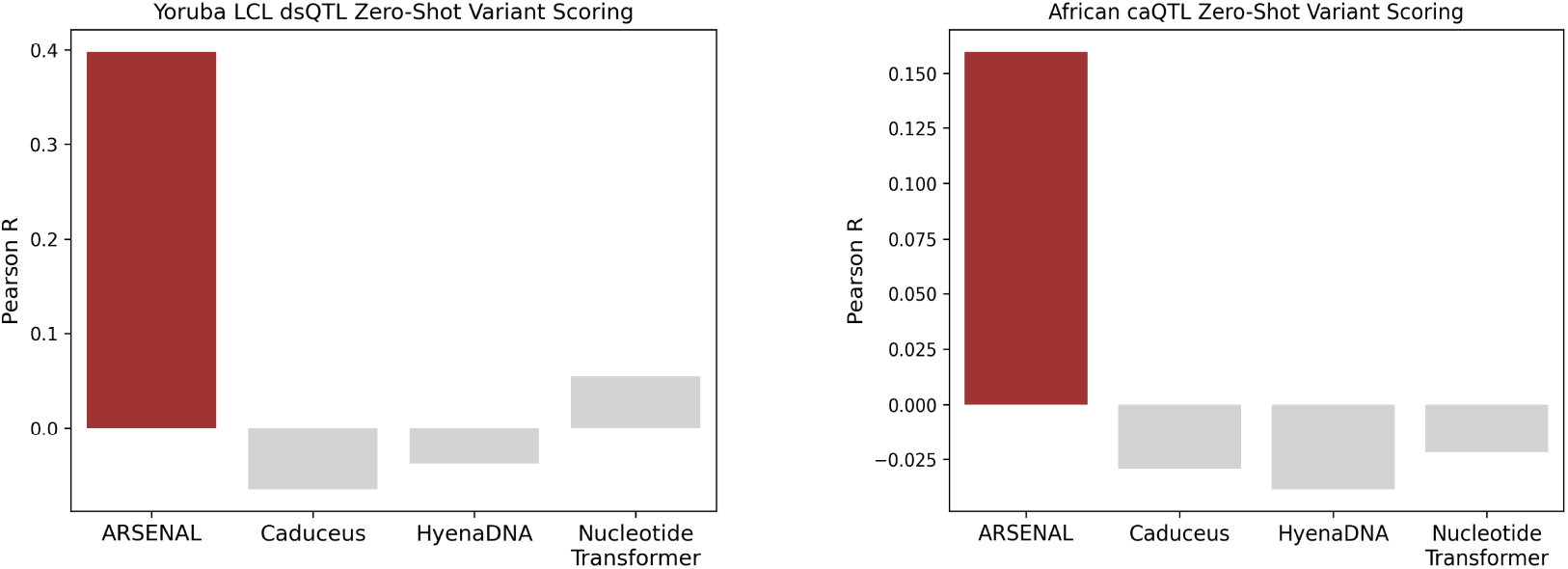
Zero-shot regulatory variant scoring on DART-EVAL QTL benchmarks. Pearson correlation between model likelihood-ratio scores and measured effect sizes for Yoruban LCL dsQTLs and African caQTLs. Scores for comparison DNALMs are from DART-EVAL [25].

### 2.4 ARSENAL embeddings improve supervised sequence models of chromatin accessibility

While likelihood-based zero-shot scoring is useful, many practical applications require supervised sequence-to-function models trained to predict cell-type-specific regulatory and transcriptional readouts from DNA sequence [4, 24, 3, 5, 18]. Such supervision is essential because regulatory mechanisms are context dependent: different cell types express distinct transcription factors and cofactors, use partially distinct motif vocabularies, and can assign different quantitative effects to the same sequence features across assays such as chromatin accessibility, TF binding, and transcription initiation. We therefore asked whether ARSENAL embeddings provide transferable regulatory representations that improve supervised prediction.

We adapted ChromBPNet, a strong short-context model of chromatin accessibility profiles (from DNase-seq or ATAC-seq experiments) [24], by replacing its one-hot input encoding with ARSENAL per-base embeddings while keeping the overall architecture and prediction task unchanged (Methods). ChromBPNet predicts peak-resolution total read counts and base-resolution distribution of read profiles over central 1,000 bp intervals using 2,114 bp of local sequence context. We trained embedding-based and one-hot baselines on DNase-seq data from five cell lines and evaluated held-out test chromosomes. Across all five cell lines, ARSENAL embeddings improve total count prediction relative to the same ChromBPNet-style architecture trained directly on one-hot sequence (Fig. 5), indicating that ARSENAL representations capture regulatory features that transfer to supervised chromatin accessibility prediction.

**Figure 5:**
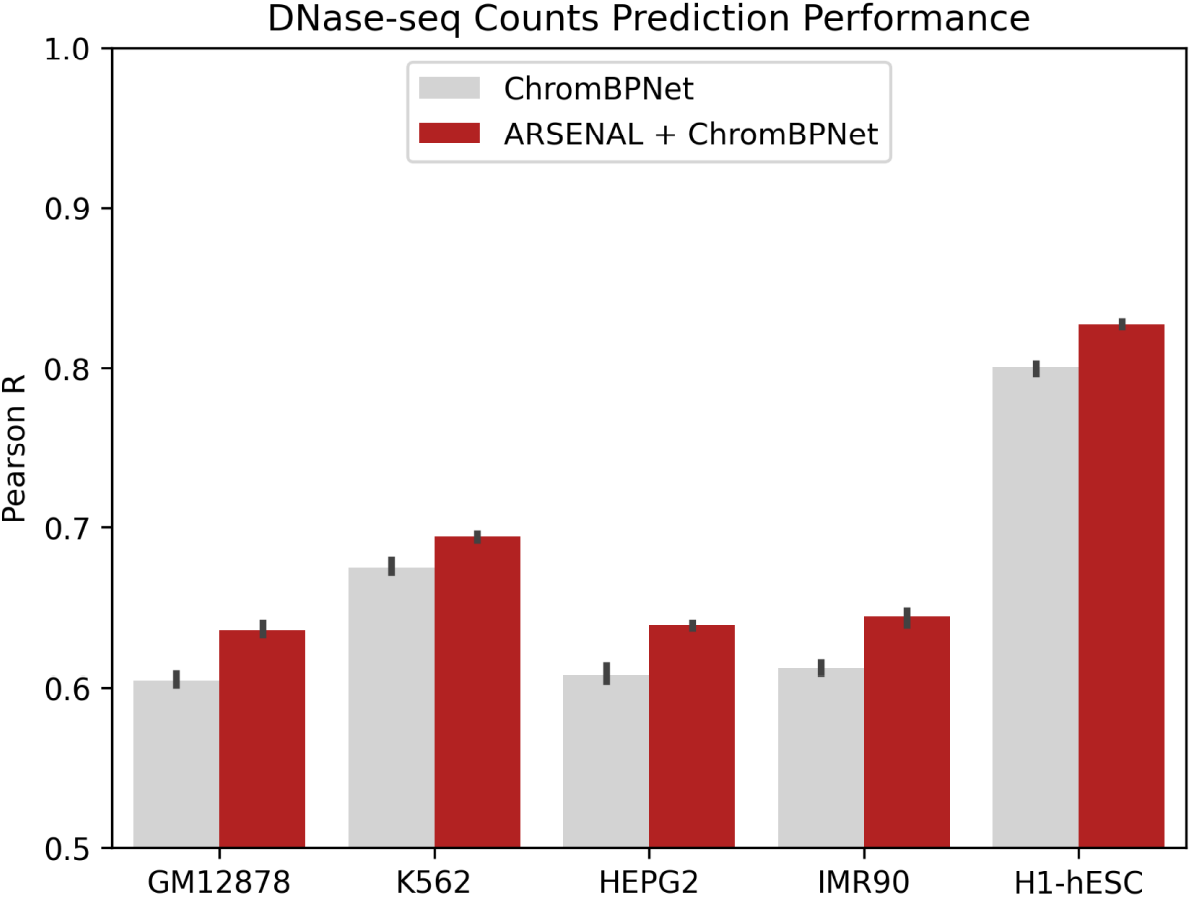
ARSENAL embeddings improve supervised chromatin accessibility prediction. Pearson correlation between predicted and observed DNase-seq total counts on held-out test chromosomes for ChromBPNet-style models trained with one-hot sequence inputs or ARSENAL embeddings across five cell lines.

We next assessed whether these gains translate to counterfactual regulatory QTL prediction. For models trained on the reference genome and DNase-seq data from the GM12878 lymphoblastoid cell line (LCL), variant scores are computed as the log fold-change in predicted accessibility between sequences containing the alternate versus reference allele. ARSENAL-enhanced ChromBPNet improves variant scoring on the LCL dsQTL and caQTL benchmarks used above (Table 1). This improvement is notable because performance on the supervised training objective does not necessarily correlate with performance on counterfactual variant effect prediction [25]. Together, these results show that ARSENAL embeddings provide biologically meaningful representations that improve both supervised regulatory prediction and downstream variant interpretation.

**Table 1:**
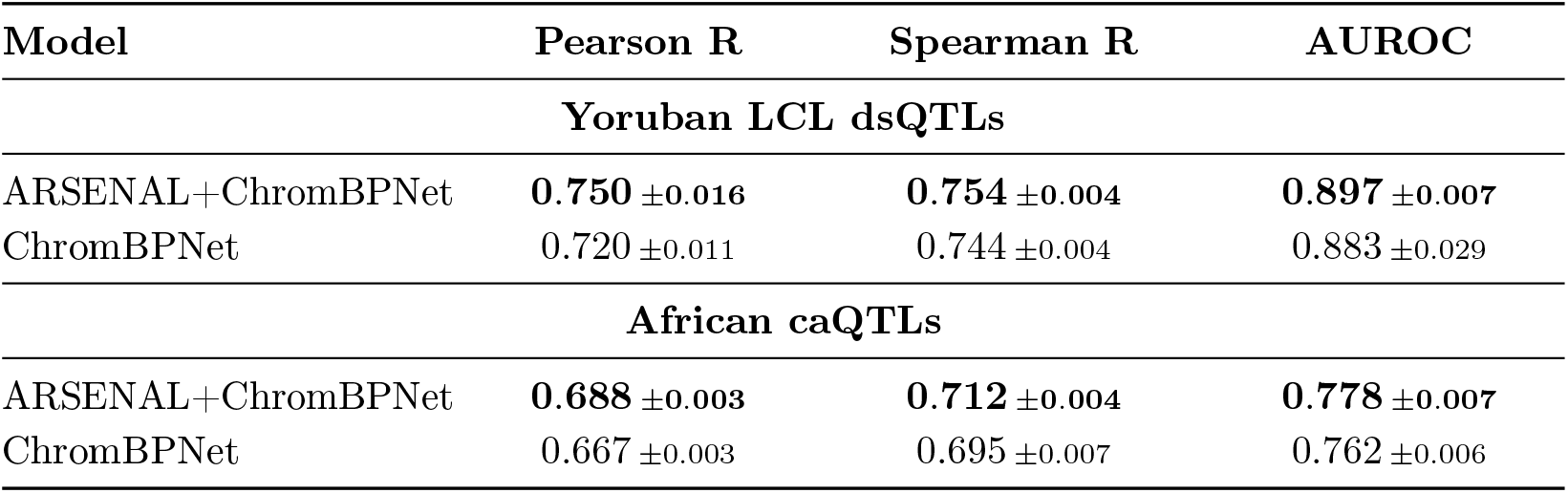
Supervised regulatory variant scoring on Yoruban LCL dsQTL and African LCL caQTL datasets. Variant scores are computed as the log fold-change in predicted accessibility between alternate and reference allele sequences using GM12878-trained models.

### 2.5 ARSENAL supports targeted regulatory sequence generation

Finally, we evaluate ARSENAL as a generative prior for regulatory sequence design. Synthetic regulatory elements are useful for therapeutic vector design, functional screening, and systematic dissection of cis-regulatory grammar [27, 12, 16, 14]. DNALMs provide a natural framework for this problem because they define a learned sequence distribution that can be sampled while maintaining similarity to regulatory DNA. The more demanding setting is *targeted* generation, where sequences must satisfy user-specified functional constraints rather than merely resemble the training distribution.

To test this capability, we combine ARSENAL with a user-defined objective and perform iterative sequence optimization using beam search. Starting from an initial sequence, we repeatedly generate candidate edits by masking and resampling a specified fraction of positions, then retain the highest-scoring candidates under the target objective. The masking rate, number of resampling steps, and temperature control the tradeoff between sequence diversity and fidelity to the starting sequence (Methods). Although this framework can support any scalar objective, here we use objectives defined by predicted chromatin accessibility from pretrained ChromBPNet models in one or more cell types.

Using this approach, we first generate sequences optimized for graded predicted accessibility in HepG2 (Fig. 6). The resulting sequences show a clear progression in predicted coverage, and ChromBPNet contribution scores indicate that higher-activity designs preferentially introduce stronger or more numerous motif instances, including CTCF motifs. We next optimize for cell-type specificity by generating sequences with differential predicted accessibility between HepG2 and H1-hESC. The optimized sequence sets show clear separation in predicted accessibility across the two cell types. TF-MoDISco analysis of ChromBPNet contribution scores reveals motif differences consistent with the design objective: HepG2-specific sequences are enriched for FOX, HNF, and C/EBP family motifs, whereas H1-hESC-specific sequences preferentially contain OCT4 and SOX2 motifs and OCT4/SOX2 motif combinations (Fig. 6C).

**Figure 6:**
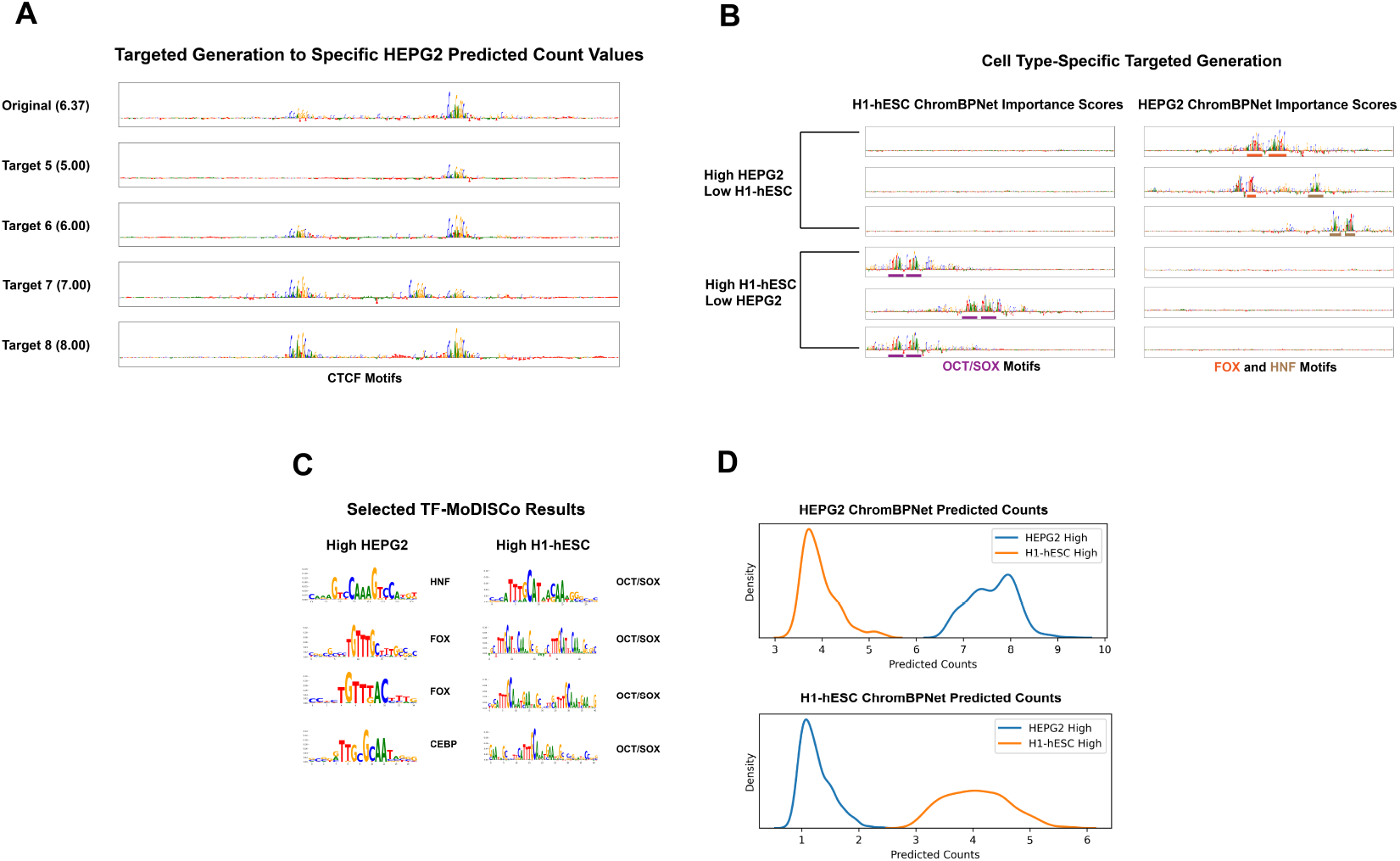
Targeted regulatory sequence generation using ARSENAL with ChromBPNet. (A) Example sequences optimized for specified predicted accessibility levels in HepG2. (B) Example sequences optimized for differential predicted activity between HepG2 and H1-hESC, with selected motifs highlighted. (C) TF-MoDISco motifs discovered from ChromBPNet contribution scores for cell-type-specific generated sequences. (D) ChromBPNet predicted count distributions for sequences optimized for differential activity.

Together, these results show that ARSENAL can support flexible predictor-guided design of regulatory DNA. Because these designs are evaluated using pretrained sequence-to-function models, experimental validation will be required to determine whether the generated sequences produce the intended activities in biological assays.

## 3 Discussion

ARSENAL demonstrates that targeted short-context self-supervision can learn biologically meaningful and transferable representations of regulatory DNA. Pretraining on ENCODE cCREs concentrates learning on regulatory elements, reducing dilution by genome-wide background sequence. Despite using only 350 bp contexts and no task-specific functional labels, ARSENAL recovers motif syntax, improves zero-shot regulatory variant scoring, transfers to supervised chromatin accessibility prediction, and supports predictor-guided regulatory sequence design.

These results support a complementary perspective to long-context, whole-genome DNALMs. Scaling context length, model size, and training corpus size can be potentially valuable, especially for modeling distal regulatory interactions or genome organization, but these factors alone do not guarantee learning of local regulatory grammar when functional signal is sparse relative to background composition. For tasks such as motif discovery, local variant scoring, supervised transfer, and short regulatory sequence design, representations that prioritize motif-scale syntax can be more useful than representations dominated by repetitive or compositional features.

Several limitations remain. ARSENAL is cell-type agnostic and does not directly model context-specific regulatory activity without downstream supervision. Its 350 bp context captures local motif grammar but not long-range dependencies such as enhancer–promoter coupling. More generally, DNALMs can learn spurious sequence patterns, including repeats, at high confidence, and sequence-only learning is constrained by the frequency and context of motifs in the training distribution. Supervised sequence models trained on cell-type-specific biochemical profiles can more directly emphasize rare but active motifs in a given cellular context. Our sequence generation experiments are also evaluated primarily *in silico* using pretrained predictors and require experimental validation.

We also evaluated a Fourier-domain likelihood regularizer inspired by prior Fourier attribution priors for supervised regulatory models [33]. In an earlier version of this work, this regularizer appeared to improve motif discovery and variant scoring, but subsequent analysis showed that the apparent gain was confounded by a floating-point precision difference between model trainings. After correcting this issue, the regularized and unregularized models performed similarly, and the main results therefore use the simpler model trained with standard MLM. Biologically motivated regularizers may still be useful for DNALMs, but they require careful specification and controlled evaluation.

Looking forward, this work suggests several directions. Expanding targeted pretraining beyond human cCREs to regulatory elements from related species could increase training diversity, although identifying comparable regulatory regions without extensive complementary genome annotations remains challenging. ARSENAL could also be combined with conservation- or MSA-based objectives, as in GPN-style models [6, 35], to integrate biochemical regulatory grammar with evolutionary constraint. More broadly, auxiliary objectives could be designed to encourage or suppress specific feature classes, such as repeat-aware penalties or priors tailored to other functional non-coding regimes including splice regions and UTRs. Finally, long-context DNALMs may benefit from initialization or modular augmentation with short-context representations, transferring robust local motif syntax while reserving long-context capacity for distal dependencies and higher-order genomic organization.

## 4 Methods

### 4.1 Pretraining Data

ARSENAL was pretrained on the ENCODE consortium’s official list of candidate *cis*-regulatory elements (cCREs) [10, 21]. This list comprises approximately 2.3 million high-confidence regulatory elements, spanning enhancers, promoters, and more ambiguous regions with regulatory signal. cCREs are identified and categorized according to sequencing experiments measuring chromatin accessibility (DNase-seq and ATAC-seq), histone mark occupancy (H3K27ac and H3K4me3 ChIP-seq), and transcription factor binding (CTCF and other TF CHIP-seq). Overall, the ENCODE cCREs represent the most extensive set of high-confidence regulatory elements curated for the human genome.

Data was split by chromosome. The validation set consisted of chromosomes 6 and 21, the test set consisted of chromosomes 5, 10, 14, 18, 20, and 22, and the rest of the genome comprised the training set.

### 4.2 Architecture and Pretraining

The ARSENAL model consists of three components:

- An embedder, which maps input tokens to input embeddings
- An encoder, which processes the sequence through several transformer layers
- A language modeling head, which maps these embeddings to token probabilities

The model’s *embedder* consists of a single embedding layer which maps the tokens A, T, G, C, N, and MASK to size-768 embeddings. The embedding for the N token is initialized as a vector of zeros and is not updated during training.

The model’s *encoder* consists of a transformer encoder architecture with 8 transformer blocks. Each block has an embedding size of 768, 8 heads, and a feed-forward dimension of 3072. RoPE is used to inject positional information in all layers.

The model’s *language modeling head* consists of a linear layer with output size 768 followed by GeLU, LayerNorm, and a final linear layer with output size 4.

The model is designed to process sequences of length 350, which is the maximum size of any cCRE. Any shorter cCRE was expanded to this length. We employed two data augmentation strategies in training. First, sequences were reverse-complemented with probability 0.5. Second, each sequence was shifted by a randomly chosen value between -50bp and 50bp.

We updated model weights based on predictions on 15% of training tokens. 80% of these were masked in the input sequence, 10% were randomly mutated, and 10% were left unchanged. We did not predict on any N’s in the input sequence.

Repeat regions represent a unique challenge for DNALMs, as language models will naturally learn them with high confidence, potentially obscuring informative regulatory features. We therefore did not update any weights based on predictions within contiguous repeat stretches of at least 30bp. These stretches were defined using lowercase letters in the reference genome. However, more work is required to determine the most optimal approach to handling these sequences.

Training took place for 150 epochs with a learning rate of 1*e*^−4^.

### 4.3 Zero-Shot Visualization and Motif Discovery

For our visualization or TF-MoDISco analyses, model likelihoods for a sequence were produced by masking each position in turn and utilizing the rest of the sequence as context to predict on the masked position. (For HyenaDNA, which is autoregressive, likelihoods were generated using a single forward pass without masking.) Likelihoods were then normalized according to the following formula, as per [15]:

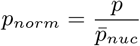

Where 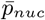 is the average predicted likelihood over the sequence for the nucleotide in question.

We performed TF-MoDISco on two datasets, both produced by the ENCODE consortium. Promoter regions were obtained by using peaks for PRO-cap, a sequencing method for transcription initiation, in the K562 cell line. We also obtained a set of DNase-seq chromatin accessibility peaks in the HEPG2 cell line. The relevant ENCODE IDs for these datasets are ENCSR261KBX and ENCSR149XIL respectively.

Peaks in these datasets are of variable length, and in all cases, we focused on the central 350bp of each region, where most motifs are located. In addition, to mitigate the presence of extraneous features in the final results, we omitted all sequences with contiguous repeat stretches of at least 50bp. Our analysis is still relatively effective without this step, but as TF-MoDISco is dependent on feature enrichment, restricting to sequences without long repeats produces the most extensive and targeted results.

TF-MoDISco was run using the `official repository`. We utilized a window size of 350 and a maximum of 1,000,000 seqlets per metacluster.

### 4.4 Nucleotide Dependency Analysis

Our nucleotide dependency analysis followed the method introduced in [32]. As stated in that paper, the dependency *e*_*i,j*_ between positions *i* and *j* in the sequence is given by:

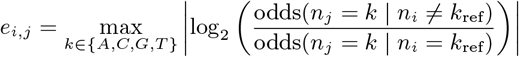

Importantly, no positions are masked when calculating nucleotide dependencies.

### 4.5 Zero-Shot Variant Effect Prediction

We utilized the zero-shot portion of the variant effect prediction task in the DART-EVAL benchmark. This task involves two datasets - a set of DNase-sensitivity QTLs from a Yoruban population, and a set of chromatin accessibility QTLs from an African population [25].

In both datasets, variants are defined according to their chromosome, position, reference allele *n*_*ref*_ , and alternate allele *n*_*alt*_. Each variant has an experimentally derived effect size, which is obtained by calculating the difference in the relevant quantity between sequences with each allele.

We score each variant by first extracting the 350 bp sequence centered upon the variant position. We then mask that position and obtain predicted likelihoods using our model. Our score *s* is then given by a simple log-likelihood ratio:

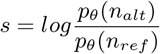

We then evaluate the quality of the variant scores by comparing to the observed effect sizes. Note that correlation metrics are only calculated on variants with significant observed effect, following the convention in the ChromBPNet paper (for which these datasets were initially curated) and in the DART-EVAL benchmark [24, 25].

### 4.6 Supervised Model Training

ChromBPNet is a dilated CNN model for chromatin accessibility prediction and variant scoring from local sequence context alone. The model accepts 2,114 bp input sequences, given as a four-channel one-hot matrix with one channel for each nucleotide. It outputs the total chromatin accessibility read counts over the central 1,000 bp along with the probability distribution of reads over that stretch.

For our model, we replace this one-hot representation with ARSENAL per-position sequence embeddings. The architecture and training scheme of our downstream ChromBPNet-style model are identical to the original except with two changes: the first convolutional layer must accept 768 input channels rather than 4, and we use a learning rate of 1*e*^−4^ rather than 1*e*^−3^.

ARSENAL embeddings for this task were derived by averaging the outputs of the last six layers of the model’s encoder. Additionally, since ARSENAL accepts an input of only 350 bp, the input sequence was split into chunks of size 350, with the central chunk comprising the center of the sequence. Embeddings for each chunk were obtained individually and then concatenated. As 2,114 is not evenly divisible by 350, embeddings for positions outside full chunks were calculated using the first/last 350 bp of the sequence.

We trained three sets of models on DNase-seq data from the GM12878, HEPG2, K562, IMR90, and H1-hESC cell lines. All training data is the same as from [25]. Note that for the fairest comparison, we trained regular ChromBPNet models using our own codebase as well. All models used predicted counts correlation as the metric for early stopping. The training, validation, and test splits comprised the same chromosomes used for pretraining.

### 4.7 Supervised Variant Effect Prediction

Variant scores for any supervised models in this study were calculated as initially described in the ChromBPNet study. We extract the 2,114 bp sequence centered at the position in question, and we create two versions: one with the reference allele at the central position, and one with the alternate allele at that position. Let us call these sequences *S*_*ref*_ and *S*_*alt*_ respectively.

For our supervised model *g*_*θ*_, we compute the model’s predicted accessibility read counts for each sequence, and we obtain our score *s* by calculating the log fold-change of the two values:

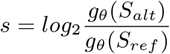

The model’s scores are then evaluated in the same manner as described for zero-shot scoring. Models for the GM12878 cell line were used for these calculations.

### 4.8 Sequence Generation

We first describe our generation loop, which is the core of our objective-guided generation workflow. This algorithm requires the definition of three main parameters:

- *N*_*i*_ - number of iterations of generation to perform
- *p*_*m*_ - probability of masking a token during an iteration
- *τ* - temperature value to scale model logits

Given a pretrained ARSENAL model *f*_*θ*_, and a seed sequence *S* the algorithm proceeds as follows:

1. For each iteration in *N*_*i*_:
  a. Create a partially masked sequence *S*_*m*_ by masking each token in *S* with probability *p*_*m*_
  b. Predict probabilities at masked positions with temperature-scaled logits: *P*_*θ*_ = SOFTMAX(*f*_*θ*_(*S*_*m*_)*/τ* )
  c. Produce a new *S* by sampling from *P*_*θ*_ at masked positions while keeping unmasked tokens the same

Given the above algorithm, we now describe our full workflow. This process requires an ARSENAL model *f*_*θ*_ and an objective function *h*, which could involve prediction using existing supervised models. The choice of *h* is flexible and task-dependent, and the parameters necessary for the function may vary.

In addition to the untargeted generation parameters *N*_*i*_, *p*_*m*_, and *τ* , we define two further parameters:

- *N*_*g*_ - number of overall generation and evaluation steps to perform
- *k* - number of sequences to retain after each generation step

Our algorithm follows a beam search strategy and is as follows:

1. Create an array *A* of generated sequences, initially with a single seed sequence *S*
2. For each step in *N*_*g*_ :
  a. For each sequence in *A*, perform untargeted generation (with parameters *N*_*i*_, *p*_*m*_, *τ* ) to produce a new set of sequences. These are added to *A*
  b. For each sequence in *A*, apply the objective function *h*
  c. If *A* contains less than *k* sequences, continue to the next iteration. If not, only retain the top *k* sequences according to the objective function

For this study, our generations involved producing sequences with certain cell type-specific properties, as defined by predictions from the relevant ChromBPNet models. Specific objectives we utilized include:

- Optimizing for one cell type: Minimizing absolute error between predicted counts and a target value in the HEPG2 cell line
- Enforcing cell-type specificity: Minimizing absolute error targeting high predicted count values in the HEPG2 cell line and low values in the H1-hESC cell line, or vice versa

Note that due to the imbalance between ChromBPNet and ARSENAL input sizes, we operated on the central 350 bp of the ChromBPNet input sequence, where most motifs are located.

We can visualize and evaluate these generations using many of the same techniques described previously. The choice of the parameters *N*_*i*_, *p*_*m*_, and *τ* determines the diversity of the generated sequences. The number of iterations and the masking probability influence the level of deviation from the original sequence, and the temperature adjusts the confidence of the model’s predictions.

For our TF-MoDISCo analyses of the cell type-specific generations, we conducted 5 generation runs each way, with 100 sequences produced per run. We then calculated contribution scores and ran TF-MoDISCo using the established protocol as found in [24].

## 5 Competing interests

A.K. is a scientific co-founder Immunera; on the scientific advisory board of SerImmune, TensorBio; is a consultant with Bristol Myers Squibb, Inari; and has a financial stake in DeepGenomics, Immunai, SerImmune, Freenome, Immunera and TensorBio.

## 6 Author contributions statement

A.P. and A.K. jointly formulated ideas, charted directions for the project, wrote and edited the manuscript.

A.P. performed all model training and experimentation.

## Acknowledgments

This work is supported in part by funds from the National Science Foundation (NSF: # 1636933 and # 1920920), NIH grant U01HG012069 and the Stanford HAI-Google Cloud Grant Program. We thank Austin Wang and Arpita Singhal for helping formulate ideas at the beginning of this project. We also thank Lei Xiong for developing a PyTorch version of ChromBPNet which was used and adapted extensively in this project.

## Appendix

### Evaluation of a Fourier-domain likelihood regularizer

In an earlier version of this manuscript, we evaluated a Fourier-domain auxiliary loss designed to encourage motif-scale likelihood structure while suppressing very short noisy features and long high-likelihood stretches. Initial ablations suggested that this objective improved performance. However, we subsequently found that the comparison was confounded by floating-point precision: the baseline model had been trained in pure bfloat16, whereas the Fourier-regularized model was trained in float32 due to PyTorch constraints. Retraining the baseline in float32 substantially improved its performance, bringing it in line with the Fourier-regularized model across the metrics evaluated here (Fig. 7, Fig. 8 and Table 2). Because the Fourier loss does not provide a clear improvement after this correction, all main-text results report the simpler MLM-trained ARSENAL model without this auxiliary loss.

**Table 2:** Comparison of supervised variant scoring performance for ARSE-NAL+ChromBPNet supervised models with and without the Fourier attribution prior. Extension of Table 1 in the main text.

| Model | Pearson R | Spearman R | AUROC |
| --- | --- | --- | --- |
| **Yoruban LCL dsQTLs** |  |  |  |
| ARSENAL+ChromBPNet<br>(With Prior) | **0.754** $\pm 0.025$ | **0.757** $\pm 0.022$ | 0.896 $\pm 0.0167$ |
| ARSENAL+ChromBPNet<br>(No Prior) | 0.750 $\pm 0.016$ | 0.754 $\pm 0.004$ | **0.897** $\pm 0.007$ |
| ChromBPNet | 0.720 $\pm 0.011$ | 0.744 $\pm 0.004$ | 0.883 $\pm 0.029$ |
| **African caQTLs** |  |  |  |
| ARSENAL+ChromBPNet<br>(With Prior) | 0.687 $\pm 0.005$ | **0.716** $\pm 0.003$ | 0.776 $\pm 0.008$ |
| ARSENAL+ChromBPNet<br>(No Prior) | **0.688** $\pm 0.003$ | 0.712 $\pm 0.004$ | **0.778** $\pm 0.007$ |
| ChromBPNet | 0.667 $\pm 0.003$ | 0.695 $\pm 0.007$ | 0.762 $\pm 0.006$ |

**Figure 7:**
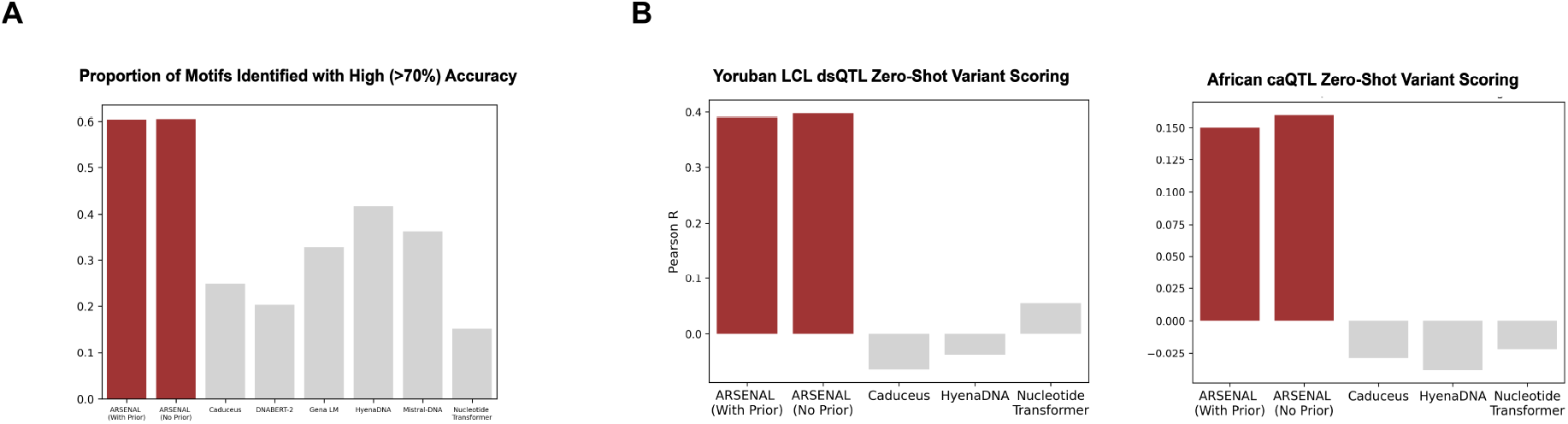
Comparison of zero-shot motif discovery and variant scoring performance for ARSENAL models with and without the Fourier-domain likelihood regularizer.

**Figure 8:**
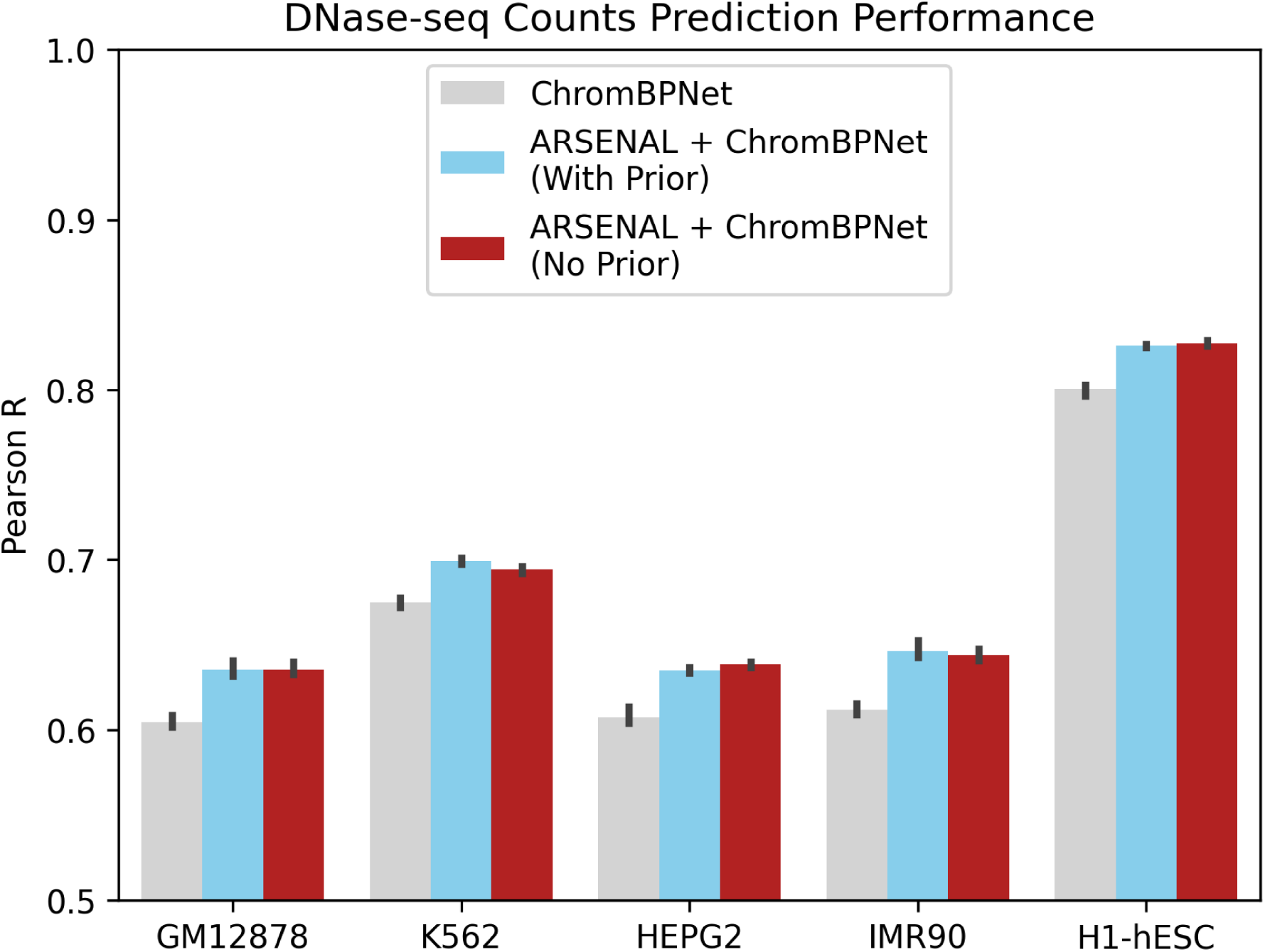
Comparison of supervised chromatin accessibility prediction performance for ARSE-NAL+ChromBPNet models with and without the Fourier-domain likelihood regularizer.

The Fourier-domain regularizer was inspired by Fourier-transform attribution priors for supervised regulatory models, which penalize high-frequency components in attribution maps to improve interpretability and motif recovery [33]. In ARSENAL, we adapted this intuition to masked language modeling by applying a frequency-domain penalty to predicted likelihood profiles rather than gradients or input attributions. The goal was to emphasize motif-length features while suppressing very short noisy features and long repetitive high-likelihood regions.

More formally, let *z* = |FFT(*f*_*θ*_(*x*))| denote the magnitude spectrum of the model’s predicted likelihood profile, restricted to positive frequencies and excluding the DC component. We then *L*_1_-normalize the spectrum:

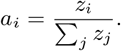

For sequence length *L*, let *C*_*H*_ = *L/*6 and *C*_*L*_ = *L/*20 denote high- and low-frequency cutoffs corresponding approximately to 6 bp and 20 bp motif scales. We assign weights to each frequency index *i*:

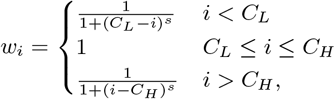

where *s* is a smoothing hyperparameter. The auxiliary loss is:

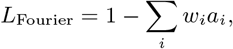

and the combined objective is:

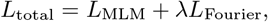

where *λ* is a loss-weight hyperparameter.

Although this regularizer does not clearly improve ARSENAL in the current setting, the result highlights an important direction for future DNALMs. Repeat regions remain a major challenge for likelihood-based sequence models, and motif recovery may depend strongly on motif frequency in the training corpus. Better-specified priors, including repeat-aware penalties or objectives that improve learning of rare but functionally important motif families, may provide more robust gains.

## Notes

### Summary of Updates

Important updates on the role of the Fourier transform-based regularizer in influencing model performance.

